# Curcumin-Sophorolipid nano-conjugate inhibits *Candida albicans* filamentation and biofilm development

**DOI:** 10.1101/2020.08.12.244186

**Authors:** Vidhyashree Rajasekar, Priti Darne, Asmita Prabhune, Richard Y. T. Kao, Adline Princy Solomon, Gordon Ramage, Lakshman Samaranayake, Prasanna Neelakantan

**Author notes:** Corresponding author: Dr. Prasanna Neelakantan, Discipline of Endodontology, Division of Restorative Dental Sciences, Faculty of Dentistry, The University of Hong Kong, Hong Kong, SAR.

## Abstract

*Candida albicans* is an opportunistic fungal pathogen that is highly resistant to contemporary antifungals, and a major reason for this appears to be their predominant, filamentation-mediated, biofilm lifestyle. Hence, agents that inhibit biofilms and filamentation of the yeast offer promise as next-generation antifungals. Curcumin is a natural polyphenol with several beneficial pharmacological attributes, yet limitations such as poor solubility, acid, and enzyme tolerance have impeded its practical utility. Sophorolipids are biologically-derived surfactants that serve as efficient carriers and delivery agents of hydrophobic molecules, such as curcumin, into biofilms. The aim of this study was to investigate the effects of a novel, curcumin-sophorolipid (CU-ASL) nano-conjugate on *Candida albicans* biofilms and filamentation. The effects of CU and ASL, in combination, and alone, were investigated on planktonic cells of the yeast. The effects of sub-inhibitory concentrations of the compounds were investigated on biofilm biomass and biofilm architecture. Their effects on filamentation was compared by scanning electron microscopic imaging, and gene expression analysis by qRT-PCR. Our results demonstrated that sub-inhibitory concentration of CU-ASL (9.37 µg/mL) significantly inhibited candidal adhesion to substrates, and subsequent biofilm development, maturation, and filamentation. This effect was associated with significant downregulation of a select group of biofilm, adhesins, and hyphal regulatory genes. In conclusion, the curcumin-sophorolipid nano-conjugate is a potent inhibitor of the two major virulence attributes of *C. albicans*, biofilm formation and filamentation, thus highlighting its promise as a putative anti-candidal agent with low toxicity and biofilm penetrative potential.

## 1. Introduction

*Candida albicans* is an opportunistic fungal pathogen, that causes mucosal, as well as life threatening infections particularly in compromised patients. Invasive, bloodstream candidal infections (candidiasis) with fatality rates as high as 40% have been reported by some [1-3]. Patients undergoing cytotoxic chemotherapy, those on long-term antibiotics, diabetics, as well as elderly denture wearers are specifically at high risk for mucosal candidal infections [1]. The ability of this yeast to form extremely tenacious biofilms on abiotic and biotic surfaces, and its morphogenic transition from yeast to hyphal/ filamentous phase, are well known as major reasons for the remarkably increased recalcitrance and tolerance of this yeast to existing antifungal agents [4].

The major drug groups currently available for the management of candidiasis are polyenes, azoles, and echinocandins [5]. However, the global emergence of resistant yeast strains to even the most effective agents has seriously impeded their clinical use, and the novel antifungal pipeline is currently, virtually dry [5, 6]. This is compounded by the fact that, being a eukaryote, pathogen-specific drug targets for anti-candidal drugs are highly limited, leading to a lag in antifungal drug development in comparison to antibiotics. An alternative, innovative approach for drug design, specifically targeting the major virulence attributes of *C. albicans*, such as biofilm formation and filamentation, is therefore urgently warranted.

The use of natural compounds for drug development, has gained immense popularity, and are advocated as safe alternatives to relatively toxic, synthetic drugs [7], [8]. Curcumin, a natural polyphenol extracted from the perennial rhizome, *Curcuma longa* (Turmeric), has long been advocated as a health conferring phytochemical due to its wide ranging pharmacological attributes such as antibacterial, antifungal, antioxidant, antitumor, and anti-inflammatory action [9]. Curcumin is also active against *C. albicans*, due to the generation of fungal-toxic, reactive oxygen species (ROS) and suppression of hyphal development [10, 11]. Yet, limiting factors such as its poor aqueous solubility and acid tolerance, low bioavailability, enzymatic degradation have impeded its clinical transition into a potent antimicrobial [12]. There is, hence, a pressing need to develop a potent drug delivery system that sustains the pharmacologic activity of curcumin, at the infective focus.

Sophorolipids are surfactants derived from non-pathogenic yeasts such as *Starmerella bombicola*, and hold great promise as vehicles for hydrophobic drug delivery [12-14]. These FDA-approved, biodegradable surfactants exist in two forms, lactonic and acidic, wherein the surfactant component, acidic sophorolipid (ASL), is a bola-amphiphile with two hydrophilic ends located on either end of a hydrophobic skeletal scaffold [13]. Lactonic sophorolipids, in general, demonstrate antifungal activity at high concentrations [15]. Recently, a curcumin-sophorolipid nano-conjugate (CU-ASL) theranostic platform was shown to tremendously improve the availability of curcumin inside bacterial biofilm and cells without bactericidal effects [12]. However, the antifungal effects of CU-ASL remains to be investigated. Based on this background, we hypothesized that low, non-toxic concentrations of curcumin-sophorolipid nano-conjugate (CU-ASL) can selectively inhibit biofilm development as well as filamentation of *C. albicans*.

## 2. Materials and methods

### 2.1 Organism, media, chemicals and culture conditions

The wild type reference strain *Candida albicans* SC5314 used in this study was obtained from the American Type Culture Collection (ATCC). The culture was maintained in Sabouraud Dextrose agar (SDA) at 37 °C in aerobic conditions. The inoculum for each experiment was prepared by taking a loop full of a single colony from the SDA plates and suspending it in Yeast Peptone Dextrose broth (YPD). The broth culture was incubated overnight at 37 °C in aerobic conditions, centrifuged at 6000rpm for 10 min and washed twice with Phosphate Buffer Saline (PBS). A standard cell suspension of 1.5×10^7^ yeast cells were prepared by optical density measurements. CU-ASL was prepared as described previously [12]. Curcumin and acidic sophorolipid were purchased from Sigma Aldrich (St. Louis, MO, USA), and carbosynth (Berkshire, UK), respectively. All experiments were performed in triplicates on three independent occasions.

### 2.2 Susceptibility of planktonic cells

The effect of CU-ASL on planktonic *C. albicans* was determined by measuring the OD_595_. Curcumin (CU) and acidic sophorolipid (ASL) were used as controls. Initially, a stock solution was prepared by dissolving CU-ASL, ASL in YPD broth, and CU in YPD broth + 10% acetone. Acetone (10%) was chosen as the solvent for CU after extensive pilot studies to optimize the solubility without affecting cell viability [16]. A working concentration of 100 µg/ml of CU, ASL, and CU-ASL was added to sterile 96-well polystyrene plates and serially diluted up to 0.585 µg/ml. The culture was prepared as mentioned earlier and 10 µl was added into each well. Untreated standard cell suspension with fresh YPD and YPD + 10% acetone were considered as control and vehicle control respectively. The plates were then incubated in aerobic conditions at 37 °C for 24 h to ascertain the minimum inhibitory concentration of the CU-ASL combination.

### 2.3 Evaluation of biofilm inhibition

A range of sub-inhibitory concentrations (sub-MIC) of the compounds was determined based on the aforementioned planktonic cell studies. The inhibition of biofilm formation by the sub-MIC concentrations was determined by measuring the biofilm biomass using the crystal violet (CV) assay [17]. Biofilm inhibition was investigated before and after the initial adhesion phase of *Candida albicans*, as described below [1], [17].

For treatment *before the initial adhesion of the yeasts*, 100 µl of varying concentrations of CU and CU-ASL were added to 10 µl of standard cell suspension. For treatment *after the initial adhesion phase*, 100 µl of the standard cell suspension was inoculated into the multiwall plates and incubated for 90 min. Next, the loosely attached cells were washed with PBS and the adherent cells were exposed to varying concentrations of 100 µl of the compounds. Untreated cell suspensions with fresh media and untreated cell suspension with 10% acetone were maintained as control and vehicle control, respectively. The plates were then incubated under aerobic conditions at 37 °C for 24 h. Afterwards, the planktonic cells were removed and biofilms were washed twice with PBS and stained with 0.1% crystal violet for 15-20 min. The residual dye on the biofilm biomass was solubilised by adding 33% v/v acetic acid, transferred to another 96-well plate, and the absorbance was measured at OD_570_ using a multimode detector.

### 2.4 Effect on filamentation

The effect of CU-ASL on filamentation was determined by allowing *C. albicans* to form hyphae on a solid substrate under hyphal inducing conditions [18]. The inoculum was prepared by culturing *C. albicans* overnight at 30 °C in YPD + 10% FBS and the OD_520_ was adjusted to yield a concentration of 1.5×10^7^ CFU/mL cells. For hyphal development, 300µL of the inoculum was added to each well along with 3 mL of varying concentrations of CU-ASL and ASL (9.37-0.585 µg/mL) prepared in fresh YPD + 10% FBS, and the plates were incubated at 37 °C for 24 h under aerobic conditions. The plates were fixed with 2.5% glutaraldehyde (Sigma Aldrich, USA) for 2 h and dehydrated in a series of ethanol solutions. Samples were then dried overnight, sputter-coated with gold and hyphal development was observed under a scanning electron microscope (SU1510, HITACHI, Minato-ku, Tokyo, Japan).

### 2.5 Confocal Laser Scanning Microscopic (CLSM) analysis

To characterize the biofilm inhibitory effects, CU-ASL was added to chamber slides (µ 8-well plate chamber slides, ibidi GmbH, Gräfelfing, Germany) with 10µL of standard cell suspension. Untreated *C. albicans* biofilms were considered as controls. The plates were then incubated in aerobic conditions for 24 h. After the incubation period, the biofilms were washed with PBS and stained with 100 µL of Live/Dead™ stain (Molecular Probes, Life Technologies, Eugene, Oregon, USA). The slides were incubated in dark for 30 min, and the stained biofilms were visualized under CLSM (Olympus FV1000, Tokyo, Japan). Z-stack images were obtained from five randomly chosen regions and the attached cells/mm^2^ were analysed using the CellC software [19, 20].

### 2.6 Effect of CU-ASL on gene expression

To elucidate the potential mechanisms by which CU-ASL inhibits *C. albicans* biofilms and filamentation, gene expression analysis was performed using qRT-PCR. Biofilms were developed with and without CU-ASL on 6-well plates and incubated under aerobic conditions for 24 h at 37 °C. PBS-washed biofilms were then scraped and collected in 1.5 mL centrifuge tubes. The samples were centrifuged at 14000×*g* for 10 min and the supernatant was discarded. The pellet was used to extract total RNA using the Promega SV Total RNA isolation system (Promega Corporation, Madison, Wisconsin, USA) following the manufacturer’s guidelines. The purity and concentration of the extracted RNA were verified using Nanodrop (Thermo Scientific, USA). RNA was then converted into cDNA using a High-Capacity cDNA Reverse Transcription Kit (Applied Biosystems, Foster City, California, USA) using the manufacturer recommended protocol. The gene expression was analyzed by qRT-PCR using 18S rRNA as the reference gene and the relative gene expression (fold change) was determined by the 2^-ΔΔCT^ method [21]. The primer sequences used in this study is tabulated (Table 1).

### 2.7 Statistical analysis

Data were statistically analysed using GraphPad Prism 8.0.2. One-way ANOVA and Dunnett’s T-test were performed to compare the significance between the control and treatment groups. The number of attached fungal cells/mm^2^ in the CLSM analysis were compared using unpaired t-test. P<0.05 was considered as statistically significant.

## 3. Results

### 3.1 Minimum inhibitory concentrations of CU-ASL, CU and ASL

A wide range of concentrations (100-0.585 µg/mL) of CU-ASL, CU and ASL were tested against the growth of *C. albicans*. CU-ASL showed more than 70% growth inhibition at 100 µg/mL, whereas CU showed ∼95% inhibition at 100 µg/mL (Figure 1). Even at lower concentrations (9.37 to 1.171 µg/mL) CU showed significant growth inhibition compared to the control, while ASL alone showed no growth inhibition at the concentrations tested. Therefore, concentrations <100 µg/mL (for CU-ASL and ASL) and <1.171 µg/mL (for CU) were considered as sub-inhibitory concentrations.

**Figure 1:**
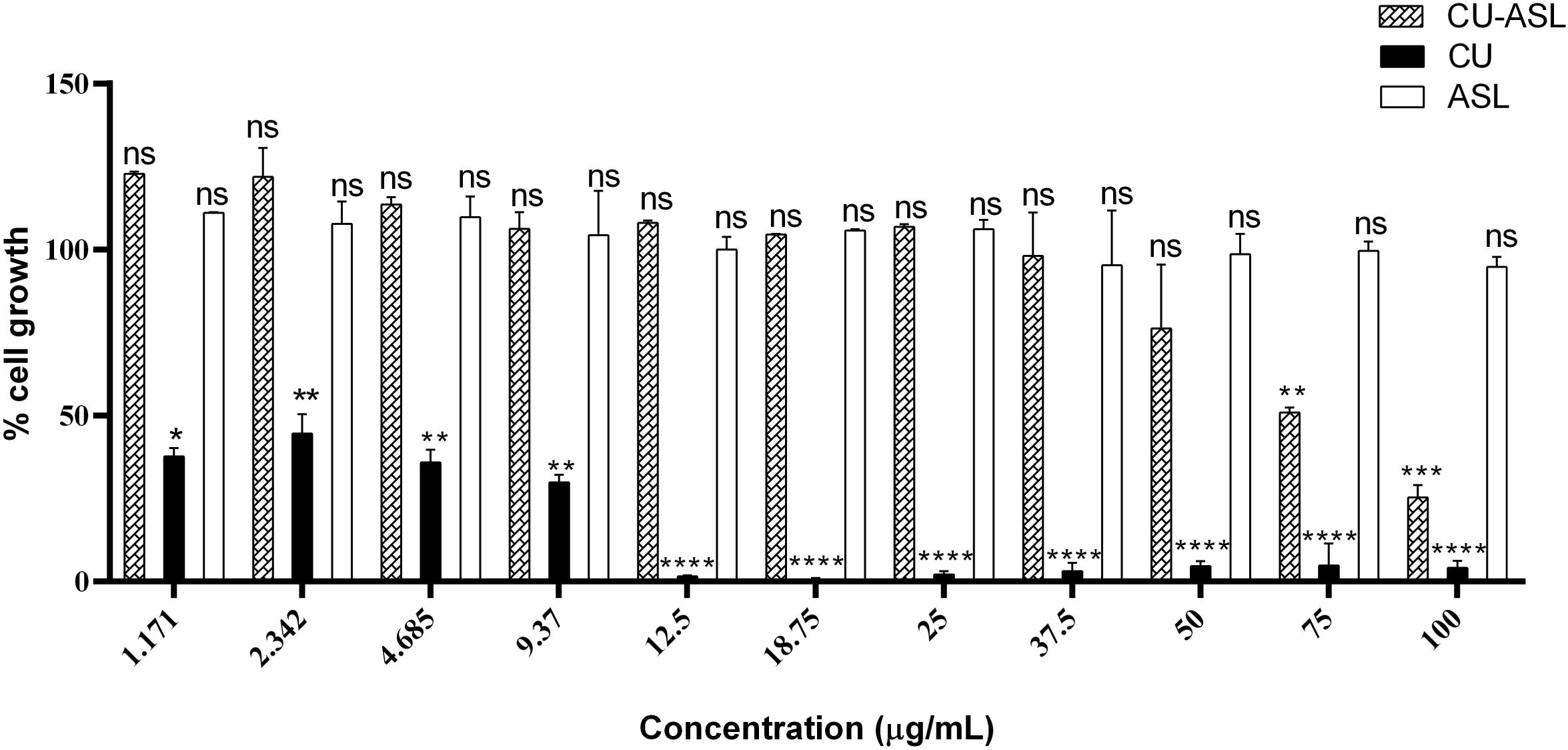
Minimum inhibitory concentrations of CU-ASL, CU and ASL. Planktonic *C. albicans* was treated with different concentrations of CU-ASL, CU and ASL. The cell growth was calculated by normalizing the inhibition data to 100%. Results represents the average of three independent experiments ± SD. *p≤0.05, **p≤0.005, ***p≤0.0005, ****p≤0.0001, ns-non-significant.

### 3.2 CU-ASL is a potent inhibitor of C. albicans biofilms in sub-inhibitory concentrations

Here, we evaluated the effect of sub-inhibitory concentrations of CU, ASL and CU-ASL on inhibition of candidal biofilm development. Based on the MIC results above, we tested a range of sub-inhibitory concentrations for biofilm inhibition. We mechanistically divided the treatment into two phases of adhesion as before and after initial adhesion of *Candida albicans* to evaluate the biofilm inhibition [1]. Within the range of sub-inhibitory concentrations tested, only concentrations below 9.37 µg/mL of CU-ASL were able to inhibit biofilms in both the pre- and post-adhesion phases (Figure 2). By contrast, sub-inhibitory concentrations of CU (<0.5 µg/mL) were unable to inhibit biofilms, and 37.5 µg/mL ASL inhibited merely 40% biofilms after the initial adhesion phase (data not shown). Since sub-inhibitory concentrations of CU were incapable of inhibiting biofilms, this group was not investigated further.

**Figure 2:**
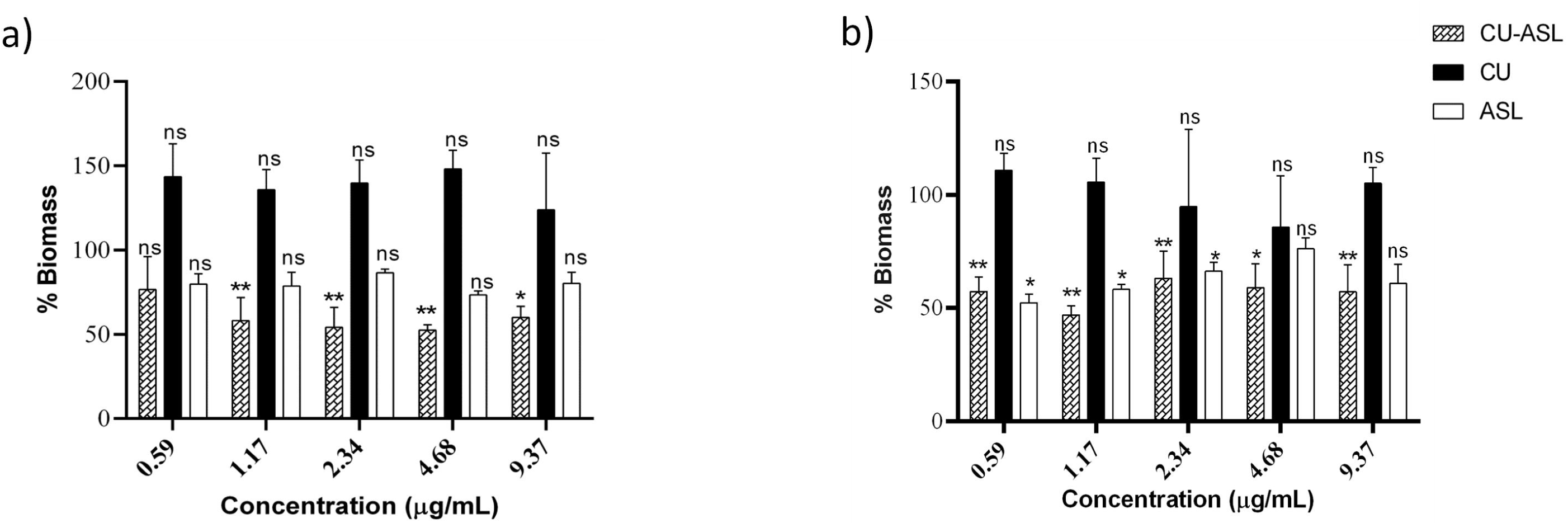
CU-ASL is a potent inhibitor of *C. albicans* biofilms in sub-inhibitory concentrations. *C. albicans* biofilms were developed in the presence of different concentrations of CU-ASL, CU and ASL treated. a) Biofilms treated at pre-adhesion phase b) Biofilms treated at post-adhesion phase. The biomass was calculated by normalizing inhibition data to 100%. Results represents the average of three independent experiments ± SD. *p≤0.05, **p≤0.005, ns-not significant.

### 3.3 Biofilm inhibitory concentrations of CU-ASL inhibit filamentation

To confirm the anti-filamentation property of CU-ASL, standard cell suspensions were grown under hyphal inducing conditions, as described previously [23]. CU-ASL abolished filamentation at 9.37 µg/mL, (Figure 3) as opposed to ASL alone that had no effect on hyphae (p <0.05). These observations reveal that sub-inhibitory concentrations of CU-ASL were able to inhibit hyphal formation more effectively than ASL.

**Figure 3:**
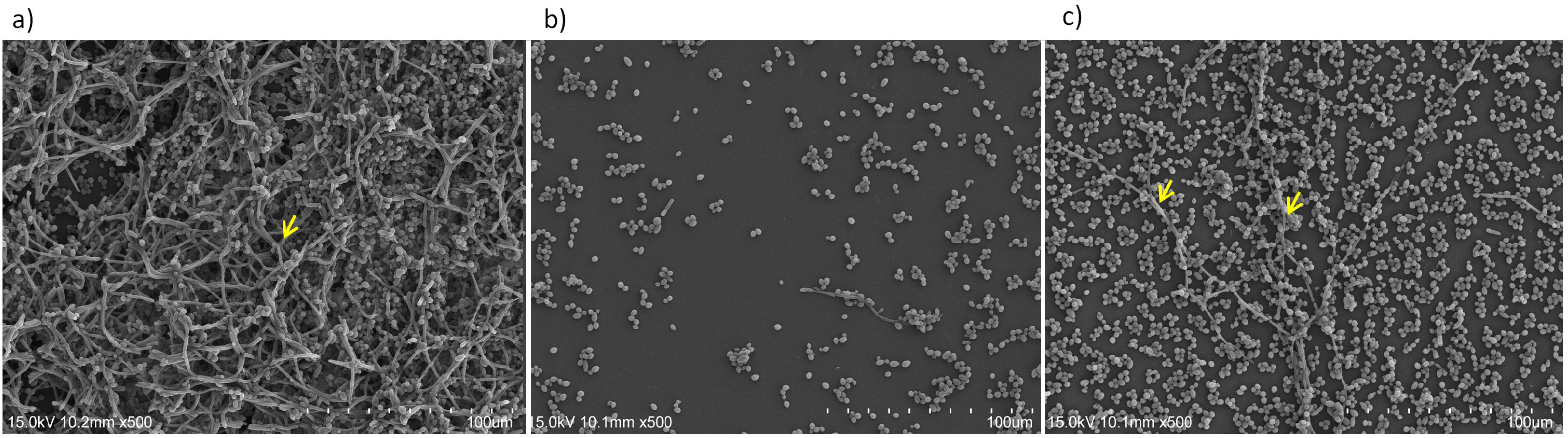
Biofilm inhibitory concentrations of CU-ASL inhibit filamentation. Scanning electron microscopic images (500x) of (a) untreated control, (b) CU-ASL (c) ASL treated *C. albicans*. The untreated control and ASL treatment show dense hyphae while CU-ASL inhibited filamentation. Yellow arrows indicate the hyphae.

### 3.4 CU-ASL treated substrates significantly impede fungal adhesion and biofilm formation

CLSM was used to investigate the architecture of CU-ASL treated biofilm and quantify the number of attached fungal cells in the substrate. The attached cells/mm^2^ was significantly reduced in CU-ASL treated biofilms when compared to the control (p<0.005; Figure 4).

**Figure 4:**
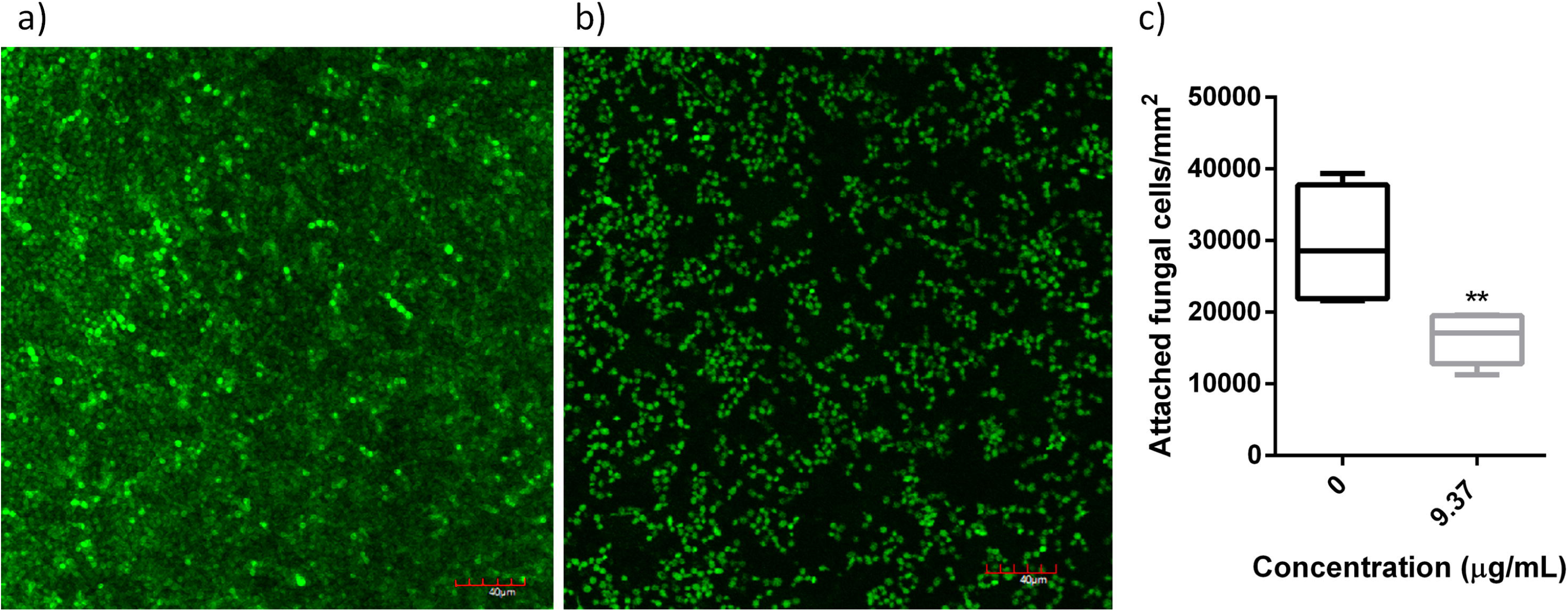
Confocal Laser Scanning Microscopic analysis of (a) control (untreated) and (b) CU-ASL treated biofilms of *C. albicans*. c) bar graph representing the number of attached live cells on the biofilm. The images were obtained at five different spots. Visualized at 40X magnification. *-p≤0.05, **-p≤0.005, ns-not significant.

### 3.5 CU-ASL downregulates multiple genes that regulate C. albicans virulence

The data shown above suggest that CU-ASL significantly inhibits biofilm and hyphal formation in *C. albicans*. Gene expression analysis revealed that the CU-ASL significantly downregulated (p<0.0001) the master regulatory genes of biofilm formation such as *ROB1, EFG1, BRG1, NDT80* and the hyphal regulatory genes *SAP4, HWP1, HYR1* (Figure 5). Specifically, *ROB1* and *EFG1* were downregulated by more than 1.5 fold. Our results also showed that the adhesin genes *ALS1, SAP8 and EAP1* were downregulated by CU-ASL treatment, substantiating the confocal microscopic observations. Amongst these, *SAP8* was downregulated by more than 2 fold.

**Figure 5:**
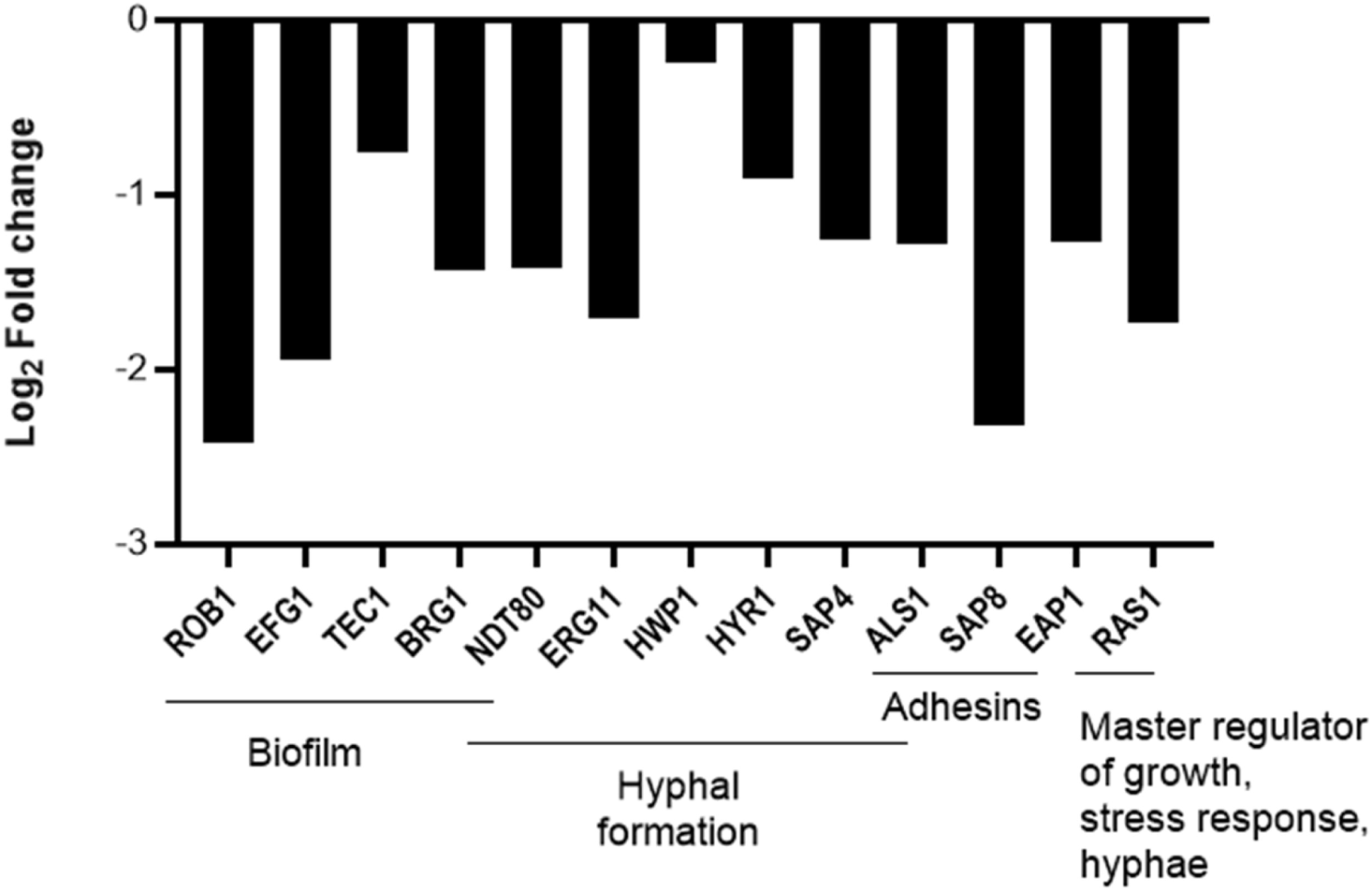
CU-ASL downregulates multiple genes that regulate biofilm formation and virulence. Differential gene expression *C. albicans* biofilm following CU-ASL treatment. Expression level of each gene is displayed after normalization with 18srRNA as housekeeping gene. The experiment was performed in duplicates. ****p<0.0001.

## 4. Discussion

Our results highlight for the first time, that the CU-ASL nano-conjugate inhibits fungal biofilm development, and retards filamentation of *C. albicans* even at very low micromolar concentrations, mainly by downregulating a battery of specific genes that regulate biofilm development and filamentation of the fungus. The antimicrobial activity of CU has been explored against a wide range of microorganisms including *C. albicans*. Alawan *et al*. [10] demonstrated that CU, ∼50 µg/mL, inhibited *C. albicans* adhesion by down regulating the adhesin genes *ALS1* and *EAP1*. Others have also shown that the hyphal repressor *TUP1* was upregulated on exposure to CU at low concentrations [9, 10, 24] Our data corroborate with the foregoing findings and in addition, demonstrate the potential anti-hyphal effect of CU-ASL at sub-inhibitory, pharmacological concentrations of the combination.

Despite such potent antifungal and anti-virulence properties of CU, its hydrophobicity renders it poorly soluble in aqueous solutions and highly unstable at physiological pH > 7 [12, 25]. It has been reported that phospholipids, liposomes and bovine serum albumin enhance the stability of CU [26]. Therefore, it is plausible that the acidic sophorolipid biosurfactant, which resembles the cell membrane by having a hydrophobic center attached to two hydrophilic tails at either end, enhances the stability and bioavailability of curcumin. Vasudevan and Prabhune [12], have described the ASL micelles as comprising three components, a hydrophilic outer shell, an aliphatic core layer and an intermediate palisade layer. While the outer shell contains sophorose and carboxylic groups, the palisade layer contains these in addition to an aliphatic region with water. This combination therefore appears to assist in actively delivering highly hydrophobic molecules such as CU into the biofilm by penetrating the extracellular polysaccharide matrix of the biofilm, though further investigations are required to test this hypothesis.

Considering the previous study on the efficient uptake of CU-ASL by bacterial cells, we hypothesized that the CU-ASL would effectively disarm the major virulence attributes of *C. albicans*, biofilm and hyphal development. Our data clearly demonstrate that this indeed is the case, as very low, non-toxic sub-MIC concentrations of the CU-ASL combination (9.37 µg/mL) had a significant suppressive effect on both candida adhesion as well as subsequent biofilm development. This phenomenon was further confirmed by qRT-PCR results, as four major transcriptional genes that govern biofilm formation, *ROB1, EFG 1, BRG 1, NDT 80* were significantly downregulated on exposure of the biofilm to the CU-ASL complex (Figure 5). Taken together, our studies (biomass analysis by the CV assay and attached cells/mm^2^ in the CLSM assay) indicate that CU-ASL has a remarkable ability to inhibit biofilms on abiotic substrates. Further studies are needed to test these effects on biotic substrates, since gene expression profiles and consequently the phenotypic effects are known to be substrate-dependent.

Filamentation associated with yeast to hyphal transition is a critical virulence attribute of *C. albicans*, that assists tissue penetration, making it the predominant human candidal pathogen. To better understand the effect of sub-inhibitory concentrations of CU-ASL on inhibition of filamentation, we performed assays on a solid substrate using SEM. From the latter observations, it was evident that CU-ASL inhibited hyphal formation at all the evaluated concentrations (data not shown), abolishing filamentation at 9.37 µg/mL (Figure 3). Interestingly, the same concentration also reduced biofilm formation in both the pre-and post-adhesion phases. On the contrary, ASL alone was unable to inhibit hyphae, except at a very high concentration of 37.5 µg/mL.

This hyphal inhibitory effect of CU-ASL could be attributed to the downregulation of the hyphal regulatory genes *ERG11, NDT80, SAP4, HWP1* and *HYR1* we observed (Figure 5). Notably, *ERG11*, which is involved in ergosterol biosynthesis pathway of lipid synthesis, has a highly polarized ergosterol-rich domain in the lipid raft of the membrane, and is responsible for filamentation. Furthermore, ergosterols are found to be associated in the septa of hypha [22]. *ERG11* is activated by the transcription factor *NDT80*. The downregulation of *ERG11* and *NDT80* also suggests that CU-ASL inhibited the ergosterol pathway, thereby reducing ergosterol polarization and hyphal formation. Hence CU-ASL exposure appears to have a wide ranging virulence suppressive effect on *C. albicans*.

A noteworthy observation of our study was the downregulation of *RAS1*, a gene involved in hyphal formation under 30°C and at low nitrogen levels. This gene is known to be a master regulator of growth, stress response and morphogenesis of *C. albicans* [27]. Although it is tempting to attribute our observation to *RAS1* suppression, at least partially, an alternative explanation could also be offered, as Hsp90, which inhibits the RAS protein, and produced under the experimental we used (i.e. a temperature of 37°C) may have contributed to this finding. Therefore, the downregulation of *RAS1* in this study may be attributed to the culture conditions rather than a true effect of CU-ASL treatment.

## 5. Conclusions

We report here for the first time, that a very low concentration of the nano-conjugate of the phytochemical curcumin, and an acidic sophorolipid in tandem (CU-ASL) is synergistically anti-fungal against an ubiquitous human pathogen *Candida albicans*, that causes common mucosal and systemic diseases. Biofilm formation and hyphal development of the yeast appear to be the major virulence attributes that are suppressed by the phytochemical-sophorolipid complex. Further studies are warranted to evaluate the antifungal effects of CU-ASL on biotic substrates including animal models.

## Acknowledgements

The authors sincerely thank Ms. Becky Cheung and Ms. Joyce Yau, Central Research Laboratories, Faculty of Dentistry, The University of Hong Kong and Dr. Yam Hill Cheung Bill, Department of Microbiology, Li Ka Shing Faculty of Medicine, The University of Hong Kong for the technical expertise and support.

## Funding

This research did not receive any specific grant from funding agencies in the public, commercial, or not-for-profit sectors.

## Competing interests

The authors declare no conflicts of interest

## Authors’ contributions

**Prasanna Neelakantan, Adline Princy Solomon, Lakshman Samaranayake**: Conceptualization; **Vidhyashree Rajasekar, Priti Darne**: Methodology, Data curation and Formal analysis; **Prasanna Neelakantan, Richard Kao**: Project administration; **Prasanna Neelakantan, Richard Kao**: Resources; **Vidhyashree Rajasekar, Prasanna Neelakantan, Lakshman Samaranayake, Gordon Ramage**: Writing - original draft; **Vidhyashree Rajasekar, Asmita Prabhune, Adline Princy Solomon, Richard Kao, Lakshman Samaranayake, Gordon Ramage, Prasanna Neelakantan**: Writing - review & editing

## Table legends

**Table 1:** Primers used in qRT-PCR analysis

## Notes

### Competing Interest Statement

The authors have declared no competing interest.

